# Molecularly imprinted nanoparticles reveal regulatory scaffolding features in Pyk2 tyrosine kinase

**DOI:** 10.1101/2023.08.05.552112

**Authors:** Tania M. Palhano Zanela, Milad Zangiabadi, Yan Zhao, Eric S. Underbakke

**Affiliations:** Roy J. Carver Department of Biochemistry, Biophysics, and Molecular Biology, Iowa State University, Ames, IA, 50011, United States; Department of Chemistry, Iowa State University, Ames, Iowa 50011, United States

## Abstract

Pyk2 is a multi-domain non-receptor tyrosine kinase that serves dual roles as signaling enzyme and scaffold. Pyk2 activation involves a multi-stage cascade of conformational rearrangements and protein interactions initiated by autophosphorylation of a linker site. Linker phosphorylation recruits Src kinase, and Src-mediated phosphorylation of the Pyk2 activation loop confers full activation. The regulation and accessibility of the initial Pyk2 autophosphorylation site remains unclear. We employed peptide-binding molecularly imprinted nanoparticles (MINPs) to probe the regulatory conformations controlling Pyk2 activation. MINPs differentiating local structure and phosphorylation state revealed that the Pyk2 autophosphorylation site is protected in the autoinhibited state. Activity profiling of Pyk2 variants implicated FERM and linker residues responsible for constraining the autophosphorylation site. MINPs targeting each Src docking site disrupt the higher-order kinase interactions critical for activation complex maturation. Ultimately, MINPs targeting key regulatory motifs establish a useful toolkit for probing successive activational stages in the higher-order Pyk2 signaling complex.

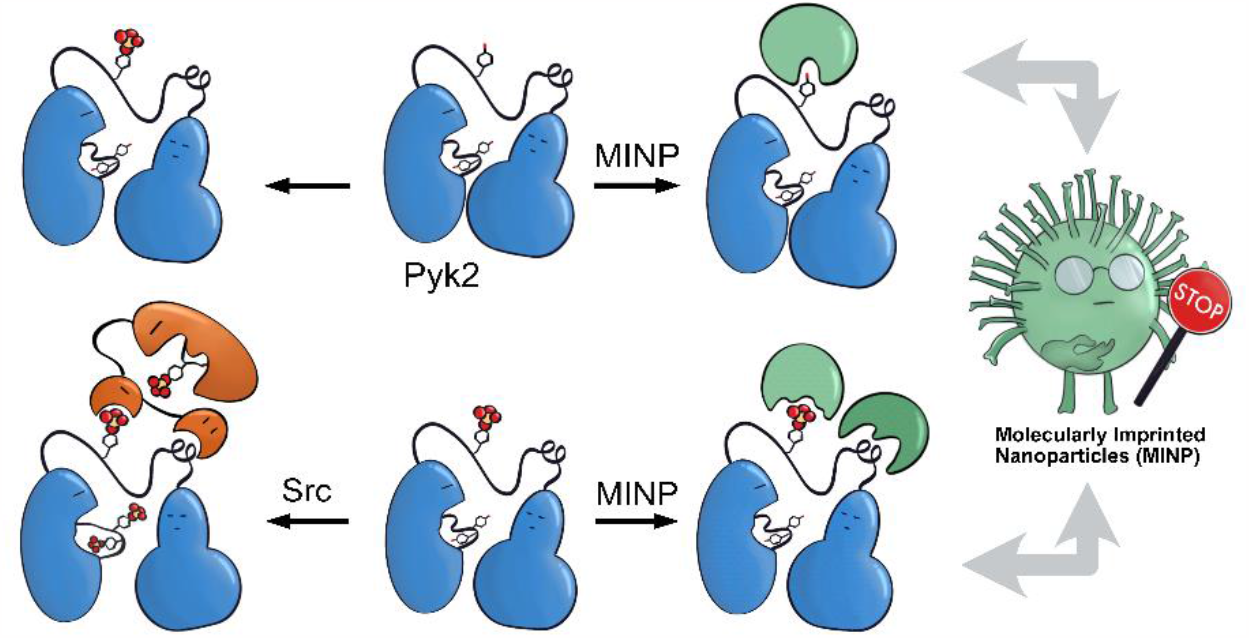

## Introduction

Proline-rich tyrosine kinase 2 (Pyk2) is a non-receptor tyrosine kinase expressed in the brain and hematopoietic cells^1^. Pyk2 signaling tunes multiple physiological processes, including cell proliferation, neuronal development, and synaptic plasticity^2^. Dysregulation of Pyk2 activity has been implicated in various pathological conditions including cancer^3^, cardiovascular diseases^4^, and neurological disorders^2^. Pyk2 shares a domain organization and significant sequence similarity (65%) with its paralog focal adhesion kinase (FAK), including an N-terminal FERM regulatory domain, a central kinase domain, and a C-terminal focal adhesion targeting (FAT) domain (Fig. 1a). Despite architectural similarities to FAK, Pyk2 has undergone evolutionary divergence, developing unique activation inputs and regulatory mechanisms. Whereas FAK is canonically activated by integrin receptor clustering at focal adhesions^5^, the cytoplasmic Pyk2 has adopted Ca2+ sensitivity in neuronal cells^1,6,7^.

**Fig. 1.**
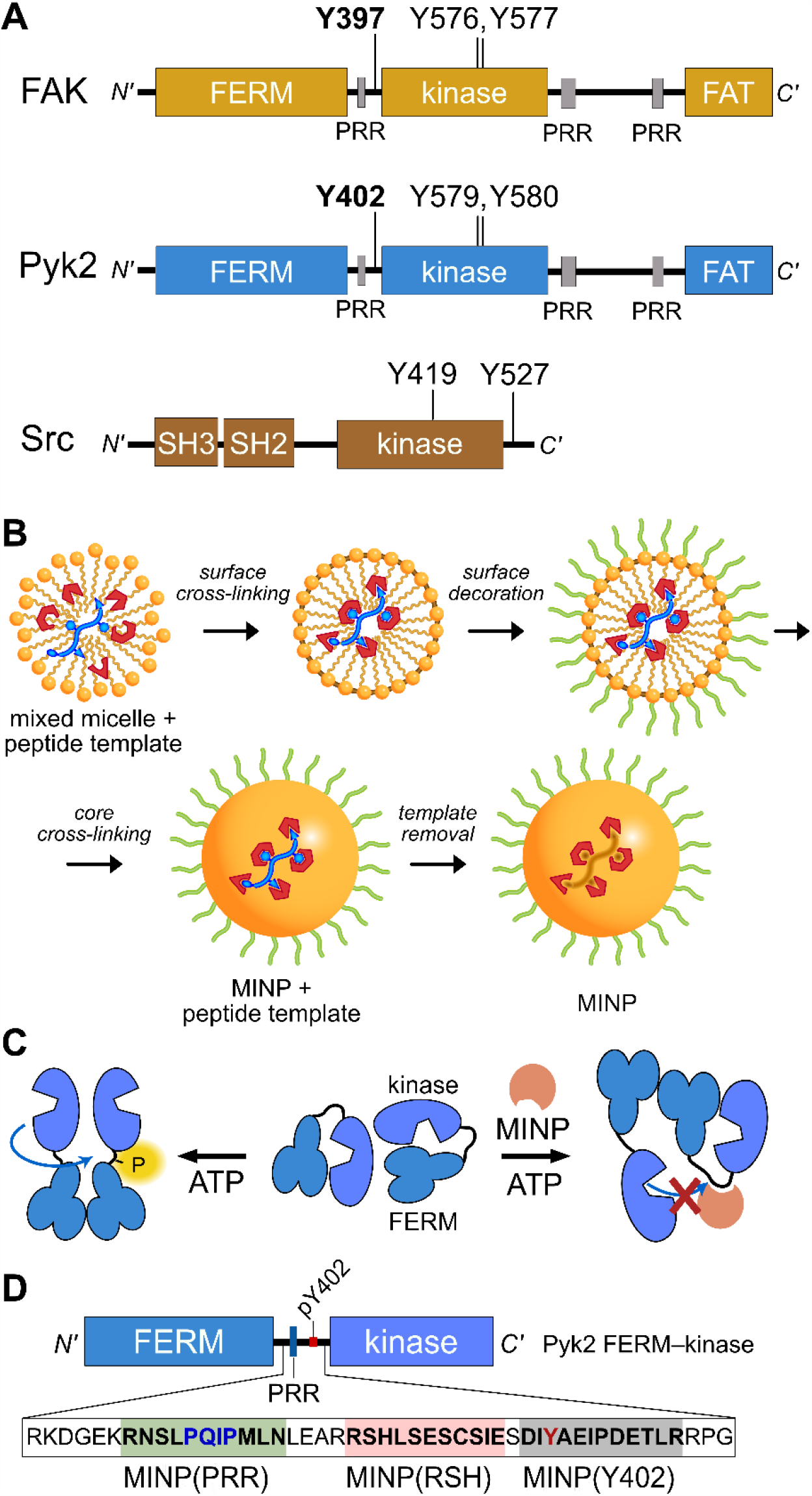
Tyrosine kinase regulatory features probed by MINPs. (a) Domain organization of Pyk2, FAK, and Src kinase. Proline-rich regions (PRR) and phosphoacceptor tyrosines are annotated with linker autophosphorylation sites in bold. (b) Cartoon representation of peptide templated MINP preparation. Mixed micelles with functional monomers are cross-linked around peptide (yellow and blue) to afford a binding pocket with shape and functional group complementarity^8^. (c) Schematic model for MINP-mediated inhibition of Pyk2 autophosphorylation. (d) Pyk2 FERM—kinase construct highlighting linker sequences used for MINP imprinting.

Despite activation differences, key regulatory mechanisms are shared between Pyk2 and FAK. Both kinases are autoinhibited by engagement of the FERM domain with the kinase C-lobe^9,10^.

The FERM domain suppresses kinase activity by obstructing access to the kinase activation loop and substrate protein docking surface. The autoinhibitory conformation is relieved by stimuli such as integrin engagement for FAK or calcium signaling for Pyk2. Stimulated conformational changes expose the kinase active site and trigger the autophosphorylation of a key tyrosine (FAK Y397, Pyk2 Y402) in the FERM–kinase linker^1,5^. The phospho-Y397 and a neighboring proline-rich region (PRR) establish a scaffold for signaling effectors like Src kinase^11,12^. The engagement of the Src SH2 and SH3 domains with the FERM–kinase linker leads to Src-mediated phosphorylation of a pair of activation loop tyrosines. Phosphorylation of the activation loop ultimately enhances the catalytic activity, which in turn promotes the phosphorylation of downstream targets.

Although the mechanistic details of Pyk2 activation are obscured by limited high-resolution structural models, investigations dissecting Pyk2 regulation revealed key features^13–15^. Pyk2 autophosphorylates residue Y402 independently of Src kinase activity. Autophosphorylation is sufficient for Src to dock and outcompete the dynamic FERM–kinase autoinhibitory interface to phosphorylate the Pyk2 activation loop tyrosines (Y579, Y580). These observations underscore the intrinsic dynamics of the Pyk2 autoinhibitory conformation and highlight the importance of the accessibility of linker residue Y402 for the maturation of the Pyk2–Src activation complex.

Despite the intrinsic capacity of Pyk2 for autophosphorylation, the regulatory mechanisms governing activation remain unclear. In FAK, the autophosphorylation site Y397 and surrounding residues are constrained in an antiparallel strand of a small FERM domain beta sheet^9^. The neighboring proline-rich region also makes intramolecular contacts with the FERM domain^16^.

Indeed, perturbations to the FAK FERM domain beta sheet (e.g., K38A) impact autophosphorylation rate^17^. To date, it is unknown whether the autophosphorylation site of Pyk2 is also constrained via secondary structure. Additionally, the regulatory contributions of the Src SH2 and SH3 domains in the Pyk2 activation complex are unclear.

To gain insight into the dynamics and signaling scaffolding of the Pyk2 autophosphorylation site, we employed chemical probes and biochemical characterizations, including peptide-binding molecularly imprinted nanoparticles (MINPs), mutagenesis, and activity profiling. MINPs are nanoparticle probes prepared from cross-linkable micelles containing template molecules such as peptides. The MINP micelles include diverse functional monomers presenting chemical moieties that provide specific interactions with template peptide features (Fig. 1b and Scheme S1). Crosslinking of the micelles generates a solid particle with docked template peptide nestled into a stable pocket lined with interaction moieties. Template removal yields cavities (i.e., imprinted sites) with shapes and interaction networks complementary to the target peptide^18,19^. The resultant “plastic antibodies” can bind peptides with high affinity (i.e., low nanomolar dissociation constants) and distinguish closely related amino acids such as leucine/isoleucine^18^, aspartate/glutamate^20^, and lysine/arginine^21^. The capacity for exquisite specificity enables MINPs to inhibit specific post-translational modifications^8,22^ or probe the functional role of a specific peptide sequence^19^. Hence, we investigated the mechanisms regulating Pyk2 activation by leveraging the distinctive features of MINPs to selectively target and probe specific Pyk2 regulatory features (Fig. 1c).

Ultimately, our study provides new insights into Pyk2 regulation by revealing that the Y402 site is conformationally constrained in the autoinhibited state due to the formation of a short betasheet with the FERM domain. Disruption of this regulatory substructure results in trans autophosphorylation of the Y402 site, allowing Src docking and subsequent Src-mediated activation loop phosphorylation of Pyk2. Our findings clarify mechanisms of Pyk2 regulation and activation while demonstrating that MINPs can serve as effective conformational probes and chemical tools to investigate protein–protein interactions.

## Results and Discussion

### The site of Pyk2 autophosphorylation is constrained via secondary structure

Previously, we reported on the efficacy of MINPs for controlling kinase activity by selective inhibition of kinase target phosphorylation sites (Fig. 1c)^23^. We demonstrated that MINPs templated with peptides representing phosphorylation target sequence motifs could site-specifically block cyclic AMP-dependent protein kinase (PKA) activity. We also examined whether MINPs could sterically block kinase access to protein phosphorylation target sites, using Pyk2 autophosphorylation as a model system. MINPs targeting the Y402 site and neighboring regions could significantly inhibit Pyk2 autophosphorylation, albeit to varying degrees depending on the specific MINP binding targets. Intriguingly, our results showed that MINP(Y402), which was designed to target the Y402 site directly, exhibited weaker inhibitory potency than MINP(PRR) and MINP(RSH) which targeted adjacent polypeptide segments (Fig. 1d). We hypothesized that the reduced inhibitory activity of MINP(Y402) may reflect lower accessibility of the autophosphorylation site in the inhibited Pyk2 FERM—kinase basal conformation^8^.

In this study, we sought to explore the correlation between MINP inhibitor potency and the accessibility of the Y402 site in the Pyk2 basal state. We first examined an AlphaFold-derived model of the autoinhibitory conformation of Pyk2^24,25^ (Fig. 2a). The AlphaFold model of Pyk2 recapitulated a beta strand sequestering the autophosphorylation site to the FERM domain as observed in the autoinhibited FAK model derived by X-ray crystallography^9^. However, the Pyk2

**Fig. 2.**
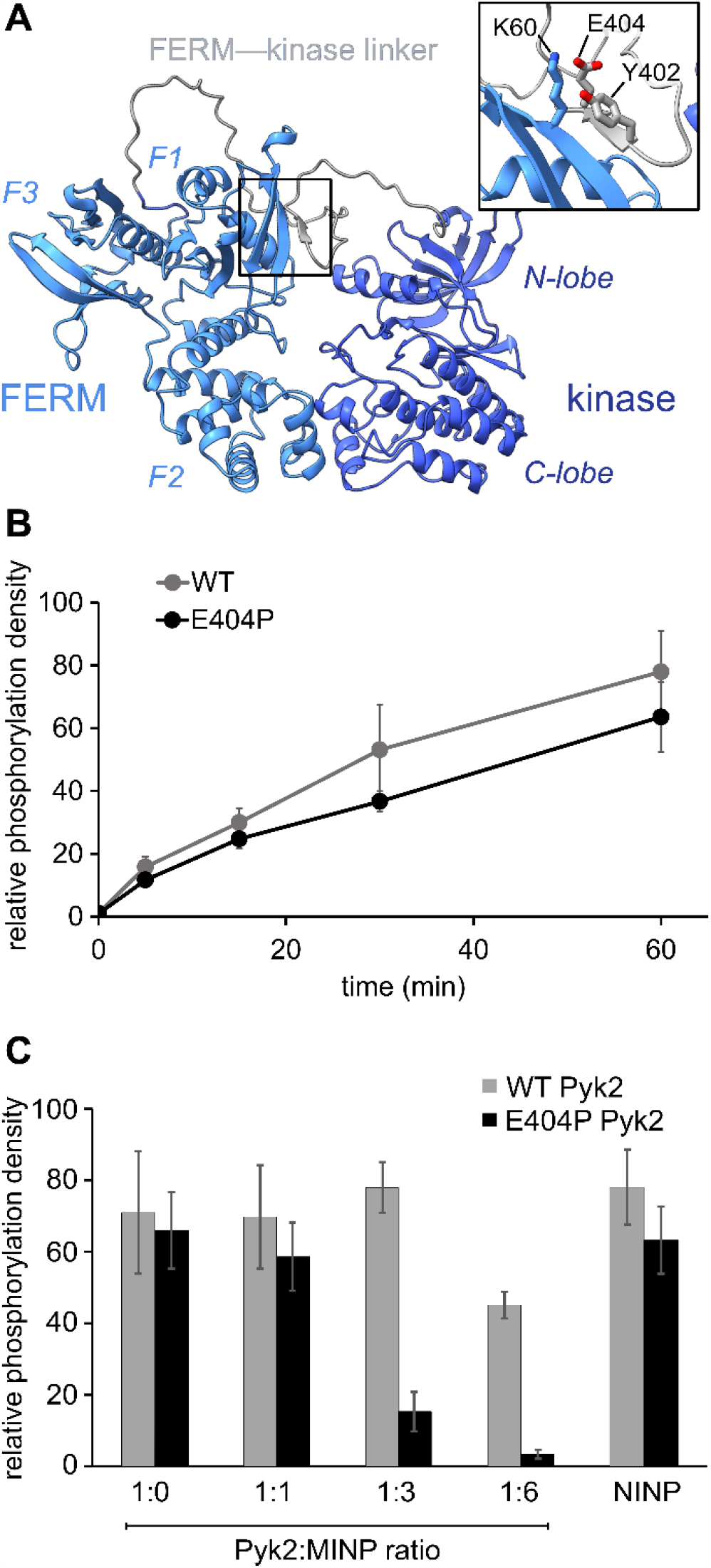
Pyk2 kinase regulatory conformations probed by molecularly imprinted nanoparticles. (A) AlphaFold-derived structural model of Pyk2 FERM—kinase with inset detailing the small β-sheet constraining Y402 in the Pyk2 linker segment colored in gray. (B) Kinase activity time course of WT and E404P Pyk2 FERM***—***kinase (0.5 μM). Tyrosine phosphorylation was detected via Western blotting with site-specific anti-phospho-Y402 primary antibody.(C)Inhibition of Pyk2 autophosphorylation by MINP(Y402) at protein:MINP ratios of 1:1, 1:3, and 1:6, with the NINP as the control at 1:6 ratio. Pyk2 concentration was fixed at 0.5 μM, and autophosphorylation was assessed 5 min after reaction initiation with ATP. Tyrosine phosphorylation was detected by dot immunoblotting with antiphosphotyrosine primary antibody (PY20). Activity assay replicates (n = 3-5) represent independent reactions with error bars signifying standard deviation.

Y402 beta strand exhibited an inverted (i.e., parallel) orientation and a very low confidence score in AlphaFold modeling. Our efforts to generate new models of autoinhibited Pyk2 using different versions of AlphaFold on a local server failed to reproduce the antiparallel, FERM-engaged beta strand observed in FAK. The apparent modeling ambiguity further motivated our investigations into autophosphorylation site accessibility in Pyk2.

Comparisons of Pyk2 AlphaFold models and the reported FAK structure enabled us to predict residues putatively responsible for stabilizing the beta sheet and sequestering the autophosphorylation site. We assessed residue-specific contributions to the stabilization of the putative interaction between the FERM domain and the FERM—kinase linker by testing a disruptive variant E404P, predicted to enhance the autophosphorylation rate. Surprisingly, we found that E404P Pyk2 variant exhibits an autophosphorylation rate similar to WT Pyk2 FERM—kinase (Fig. 2b and S1). We considered two possible explanations for this observation.

Pyk2 regulation may differ from FAK, and Y402 sequestration does not limit basal autophosphorylation. Alternatively, the local structure constraining Y402 may be stabilized by multiple cooperative interactions and/or the E404P substitution is insufficient to disrupt and liberate the linker. Importantly, kinase specificity for the local autophosphorylation motif remained a factor, and further residue substitutions risked perturbating recognition by the kinase active site.

To further interrogate the regulatory role of autophosphorylation site sequestration, we tested the inhibitory effects of site-specific MINPs. We assessed whether the E404P variant would present a more accessible binding site for MINP(Y402) and thus enhance inhibitory potency (Fig. 2c and S2). *In vitro* kinase assays of the WT and E404P Pyk2 FERM—kinase revealed that MINP(Y402) inhibited the autophosphorylation of the E404P Pyk2 variant far more effectively than WT Pyk2. Specifically, at a Pyk2:MINP ratio of 1:6, we observed near-complete inhibition of autophosphorylation of the E404P variant of Pyk2, while the wild-type (WT) Pyk2 phosphorylation was inhibited by only 30%. A non-imprinted nanoparticle (NINP), prepared without the template, exhibited negligible inhibition. The enhanced inhibitory potency of MINP(Y402) towards the E404P Pyk2 variant supports the hypothesis that the Y402 site and surrounding residues are constrained in a β-sheet conformation in the Pyk2 basal state. Thus, despite the sequence mismatch generated by the E404P Pyk2 mutation, the local structural perturbation in the Pyk2 FERM—kinase linker affords a better binding site for MINP(Y402). Based on the observed differential impact of MINP inhibition between the linker targets, we inferred that MINPs can differentiate between local polypeptide accessibility and/or conformations. Notably, the β-strand conformation is uniquely resistant to MINP recognition due to its fully extended torsion angles that differ significantly from the free, flexible peptide template imprinted on the MINP. This observation suggests that the MINP templating strategy can be leveraged to differentiate local polypeptide conformation or accessibility.

### Conformational context modulates Pyk2 autophosphorylation via the FERM domain

Our investigation into the regulation of Pyk2 autophosphorylation by MINPs revealed a reduced accessibility of the Y402 site within the inhibited conformation of Pyk2 FERM—kinase, presumably due to secondary structure engagement with the FERM domain. To further explore factors stabilizing Y402 sequestration, we returned to the AlphaFold-derived model of Pyk2 and reported autoinhibitory conformation of FAK (Fig. 2a). The modeled Pyk2 FERM—kinase indicates an electrostatic sidechain interaction between FERM F1 subdomain residue K60 and linker residue E404. Notably, E404 can interact with K60 in either antiparallel or parallel beta strand orientation. To test the contribution of Pyk2 K60 residue in stabilizing the interactions constraining Y402, we engineered alanine (K60A) and proline (K60P) variants of Pyk2 FERM—kinase construct. We investigated the impact of K60 variants on Pyk2 autoinhibition by monitoring autophosphorylation over a 30-minute time course (Fig. 3a and S3). Our results revealed that the K60A variant exhibited a >5-fold increase in autophosphorylation compared to WT Pyk2. Strikingly, introducing a proline at the K60 position led to an even higher increase in Pyk2 autophosphorylation (>10-fold increase), implying a considerable increase in linker accessibility.

**Fig. 3.**
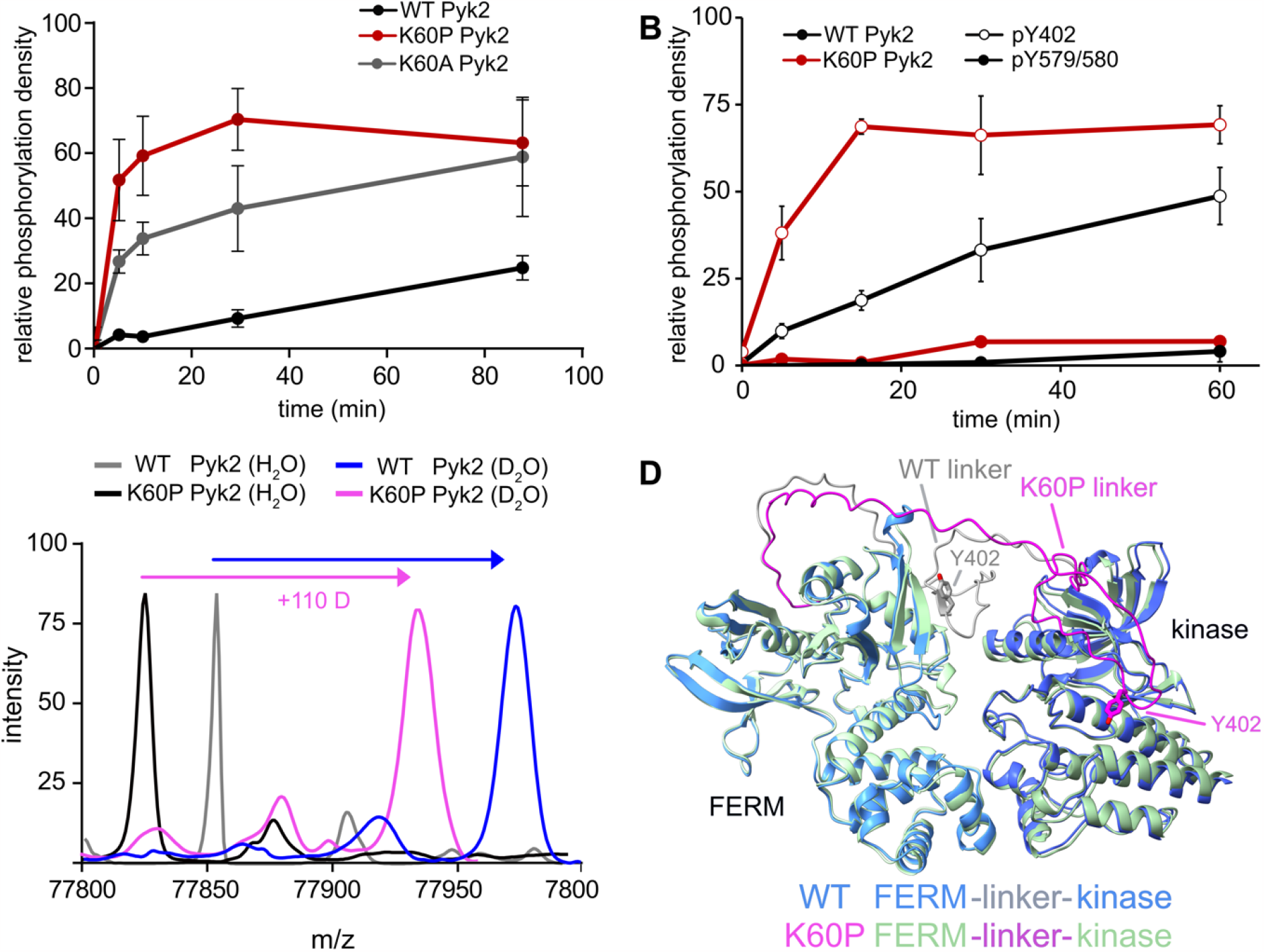
FERM domain variants modulate Pyk2 autophosphorylation. (A) Kinase activity time courses of WT Pyk2 FERM***—***kinase (0.5 μM) and variants K60P and K60A. Tyrosine phosphorylation was detected via Western blotting with pan-specific anti-phosphotyrosine primary antibody (PY20). (B) Site-specific phosphorylation was monitored by blotting with anti-phospho-pY579/pY580 or anti-phospho-Y402 primary antibody. Phosphorylation levels were quantified by densitometry. Activity assay replicates (n = 3) represent independent reactions with error bars signifying standard deviation. (C) Global HDX-MS spectra of WT and K60P Pyk2 FERM–kinase. Spectra include unlabeled (H_2_O) WT (grey) and K60P (black) superimposed with 2.5 min D_2_O exposure of WT (blue) and K60P Pyk2 (magenta). (D)AlphaFold (ver. 2.3.1) generated models of WT (blue) and K60P (green) Pyk2 autoinhibitory conformation. The FERM***—***kinase linker is highlighted with the Y402 site for WT (grey) and K60P (magenta).

The canonical mechanism for full activation of autophosphorylated Pyk2 involves Src docking and subsequent activation loop phosphorylation at Pyk2 residues Y579 and Y58014,15,26. Given the pronounced increase in Pyk2 autophosphorylation upon disruption of the putative FERM:linker interface, we tested whether a K60 variant also impacted the phosphorylation status of the activation loop. We monitored site-specific phosphorylation in WT and K60P Pyk2 FERM—kinase. Blotting with site-specific antibodies revealed that the autophosphorylation target of the K60P variant was primarily the FERM—kinase linker residue Y402 (Fig. 3b and S4). Like WT Pyk2, the K60P variant exhibited negligible autophosphorylation of its own activation loop.

Our results show that variants designed to disrupt FERM—kinase linker sequestration can increase MINP access or promote autophosphorylation. We surmised that the disruptive variants increase conformational accessibility of the linker autophosphorylation site. However, perturbing remote FERM:linker contacts could also destabilize the autoinhibitory interface between FERM F2 and kinase C-lobe subdomains (Fig. 2a)^9,10^. Indeed, previous studies established that disruption of the Pyk2 FERM F2:kinase C-lobe interface leads to increased autophosphorylation^15^. Hence, we sought to test whether the K60P mutation disrupts the global autoinhibitory conformation of Pyk2. We employed global hydrogen/deuterium exchange mass spectrometry (HDX-MS) to monitor the general uptake of deuterium throughout the intact protein. Our results revealed negligible differences in global exchange between WT and K60P Pyk2 FERM—kinase (Fig. 3c). In contrast, targeted disruption of the FERM F2:kinase C-lobe autoinhibitory interface has been shown to increase global H/D exchange by 20 – 40 deuterons^10^. Taken together, we conclude that point mutations in the FERM domain (e.g., K60P Pyk2) disrupt main chain H-bonding and ion pairing, liberating the linker segment harboring autophosphorylation site Y402. Interestingly, when challenged to predict the architecture of the K60P variant of Pyk2, AlphaFold consistently generates models with a disengaged, conformationally flexible FERM—kinase linker (Fig. 3d).

### MINP binding competes with tandem Src motifs for Pyk2—Src complex formation

In addition to kinase activity, Pyk2 signaling involves scaffolding roles, primarily attributable to the tandem proline-rich region and tyrosine phosphorylation site in the FERM—kinase linker (Fig. 1a). Pyk2 scaffolds Src tyrosine kinase which is primarily responsible for Pyk2 activation loop phosphorylation at sites Y579 and Y58015 (Fig. S5). The Pyk2—Src activation complex serves as the arbiter of downstream Pyk2 signaling cascades. The Pyk2—Src interaction is essential for the regulation of cell migration 27and modulating the structural plasticity of synapses in neuronal cells28. The binding of Pyk2 and Src is a highly regulated process that involves multiple protein domains and phosphorylation. Due to similarity with FAK, the docking of Src to Pyk2 is thought to be mediated by the binding of tandem Src SH3 and SH2 domains to Pyk2 proline-rich region (PRR) and phosphorylated Y402, respectively. The engagement of Src SH2 and SH3 domains with the Pyk2 FERM—kinase linker is thought to be a prerequisite to disengagement of the autoinhibitory conformation and phosphorylation of the Pyk2 activation loop. We validated the importance of Src phosphotyrosine engagement by testing kinase constructs missing scaffolding (Pyk2 phospho-Y402) or docking (Src SH2) features. Precluding Pyk2 autophosphorylation with Y402F substitution impairs Src-mediated phosphorylation of the Pyk2 activation loop residues Y579 and Y580 (Fig. 4a,b and S6). Likewise, Src lacking the SH2 domain (ΔSH-2 Src) is deficient in Pyk2 activation loop phosphorylation (Fig. 4b and S6d), in accordance with cell-based studies14. Taken together, Pyk2 activation loop phosphorylation requires Src docking via phosphorylated Y402.

**Fig. 4.**
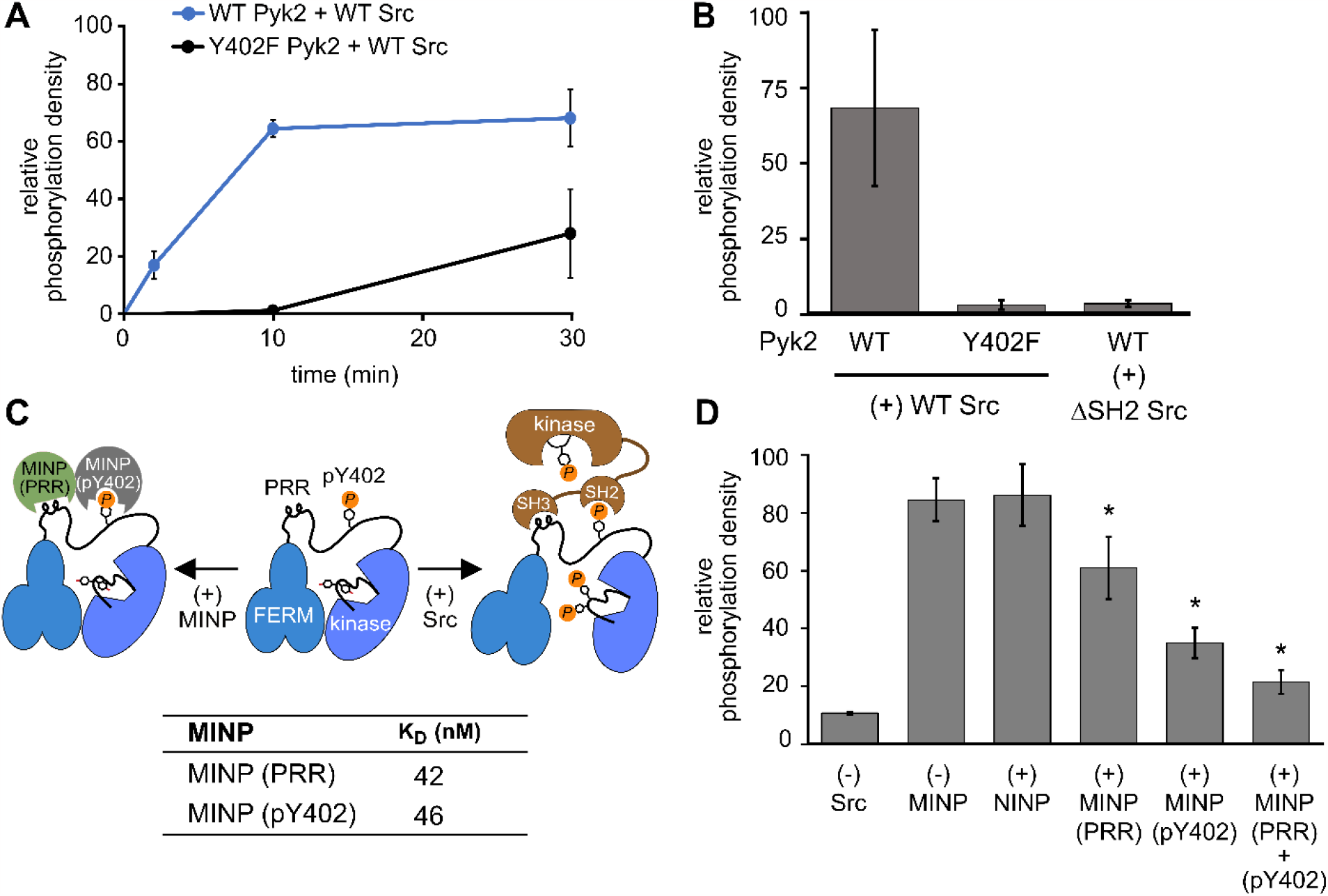
Scaffolding sites in the Pyk2—Src complex probed by MINPs. (A) Kinase activity time course of WT or Y402F Pyk2 FERM**-**kinase (0.5 μM) in the presence of Src (0.5 μM). Site-specific phosphorylation was detected by blotting with anti-phospho-pY579/pY580 primary antibody. (B) WT or Y402F Pyk2 FERM**-**kinase (1 μM) was preincubated with ATP for 10 min. Activation loop phosphorylation (pY579/pY580) was measured following addition of WT Src or ΔSH2 Src [1-142,166-536] (0.5 μM). (C) Schematic illustrating MINP-mediated inhibition of Pyk2 activation loop phosphorylation. Dissociation constants were measured via isothermal titration calorimetry for MINP(pY402) and MINP(PRR) with the corresponding peptide template. (D) Activation loop phosphorylation (Y579/Y580) of phospho-Y402 Pyk2 FERM**-**kinase (1 μM) preincubated with MINP. Src (0.5 μM) was added 3 min after MINP equilibration. Reactions were quenched 5 min after Src addition. Activation loop phosphorylation was assessed by blotting with anti-phospho-Y579/Y580 primary antibody and quantified by densitometry. Activity assay replicates (n=3) represent independent reactions with error bars representing standard deviation. *p<0.05 vs the (+) NIMP treatment.

Interested in targeted strategies to interfere with specific protein–protein signaling interactions29, we investigated the potential of MINPs to selectively target Pyk2 docking sites and disrupt the formation of the Pyk2**-**Src activation complex. Specifically, we hypothesized that MINPs imprinted with either phospho-Y402 sequence or PRR sequence (Fig. 4c) could outcompete the tandem Src motifs for Pyk2 binding. For this purpose, we assessed the inhibition of Src-mediated phosphorylation of Pyk2 activation loop tyrosines (Y579/Y580) in the presence of MINP(pY402) and MINP(PRR) alone or in combination. First, WT Pyk2 FERM**-**kinase was preincubated with ATP to allow for basal autophosphorylation to fully phosphorylate the Y402 site. Phospho-Y402 Pyk2 was pre-incubated with MINP or NINP followed by addition of Src.

The impact of MINPs on Src-mediate Pyk2 activation loop phosphorylation was monitored using site-specific activation loop anti-phosphotyrosine (pY579/Y580) antibodies. Our results reveal that blocking either the Src SH3 or SH2 domain binding site in Pyk2 leads to substantial inhibition of Src-mediated phosphorylation of Pyk2 activation loop tyrosines (Fig. 4e and S7).

To control for the possibility that MINPs interfere with Src directly rather than blocking the targeted Pyk2 motifs, we tested Src activity in the presence of MINPs and NINP. Intrinsic Src kinase activity was unaffected by MINPs (Fig. S8). Notably, despite similarly high binding affinities (Fig. 4c and S9), MINP(pY402) exhibited more potent inhibition of Src activity than MINP(PRR). The combination of MINP(pY402) and MINP(PRR), however, resulted in further inhibition of Src-mediated phosphorylation of Pyk2 activation loop tyrosines (Fig 4e). Thus, motif-specific MINP inhibition suggests that Src SH2 recognition of Pyk2 phosho-Y402 plays a critical role in nucleating the Src—Pyk2 activation complex. Nevertheless, the significant inhibition by MINP(PRR) reveals that blocking access to the Pyk2 PRR is sufficient for disruption of Src-mediated Pyk2 activation. Our results demonstrate that MINPs targeting either Src-binding motifs in Pyk2 can successfully outcompete Src domains and inhibit Src-mediated Pyk2 activation loop phosphorylation.

We also note an intriguing contrast between the activities of MINP(Y402), targeting Pyk2 autophosphorylation, and MINP(pY402), targeting Src SH2 docking via the same site modified by phosphorylation. MINP(Y402) inhibition is muted by the apparent inaccessibility of the unphosphorylated Y402 region (Fig. 2c). MINP(pY402), however, readily recognized the phosphotyrosine linker motif and potently inhibited Src activity. Given the role of ion pairing in stabilizing the sequestration of the Y402 region (Fig. 3), we speculate that Y402 phosphorylation prevents engagement with the FERM β-sheet due to local conformational changes associated with the additional negative charge. The increased accessibility of the linker upon Y402 phosphorylation may be important for Src SH2 docking.

## Conclusions

Ultimately, this study demonstrates the utility of peptide-binding MINPs as conformational probes to elucidate the intricate mechanisms regulating Pyk2 activation and Src scaffolding. By selectively targeting regulatory motifs in specific conformations, MINPs revealed how Pyk2 autophosphorylation at the long linker connecting FERM and kinase domains is restricted via interactions with the FERM domain. Targeted mutations at the linker sequestration site are sufficient to liberate the linker for autophosphorylation without disrupting the primary autoinhibitory interface between FERM and kinase. MINPs also dissected the contributions of tandem Src docking elements in the interdomain linker of Pyk2. These findings provide important molecular insights into Pyk2 regulation as both a signaling enzyme and scaffold for a higher-order kinase complex. The selective blockage of unique Pyk2 regulatory features by tailored MINPs highlights the potential for deploying MINPs as surgical inhibitors of specific signaling subsystems. Indeed, both FAK and Pyk2 are archetypal signaling hubs that serve multiple roles as signaling enzymes and organizational scaffolds of multiprotein signaling complexes^26^. More broadly, this work highlights the potential for using tailored MINPs as specific inhibitors or sensors to dissect signaling mechanisms, protein interactions, and conformational dynamics. The capacity to selectively recognize local protein conformation and post-translational modification allows MINPs to probe the molecular mechanisms of signaling function in complex systems.

## Author Contributions

T.M.P.Z, Y.Z., and E.S.U. conceived of the research; T.M.P.Z. performed experiments; M.Z. synthesized and characterized MINPs; T.M.P.Z, M.Z. analyzed data; T.M.P.Z, Y.Z., and E.S.U. wrote the manuscript.

## Acknowledgements

We thank Joel Nott of the Iowa State University Protein Facility for mass spectrometry support and Dipali Sashital for AlphaFold modeling guidance. This research was funded by the National

Science Foundation, Division of Molecular and Cellular Biosciences grant 1715411 and Division of Materials Research grants 2308625 and 2002659.

