## Supplemental Information file for "Molecularly imprinted nanoparticles reveal regulatory scaffolding features in Pyk2 tyrosine kinase"

### Methods

#### Preparation and Imprinting of Functionalized MINPs

Preparation of the functionalized MINPs was carried out as previously reported[17]. For nomenclature clarity, MINP11a and MINP11c from Ref. 17 were renamed MINP(PRR) and MINP(Y402), respectively, in this study. MINP(pY402) was synthesized for this study using a phosphorylated peptide (DIpYAEIPDETLR) mimicking the linker sequence surrounding the Pyk2 autophosphorylation site Y402. The MINP(pY402) template peptide was phosphorylated by a 3 h, 37 °C incubation of peptide with 1 μM Pyk2 FERM−kinase in buffer consisting of 50 mM HEPES pH 7.4, 150 mM NaCl, 8 mM $MgCl_2$, 5% glycerol, and 4 mM ATP. Peptide phosphorylation was confirmed via liquid chromatography-mass spectrometry (LC-MS).

MINPs were prepared with functional monomers using the following stoichiometries in the formulation: 1.5:1 for **1**/carboxylate or phosphate on the peptide template, 1:1 for **2**/arginine, and 1:1 for **3**/amine (Scheme S1). Compound **1** in methanol (37.5 μL of a 5.5 mg/mL solution, 0.6 μmol), **2** in methanol (50 μL of a 9.5 mg/mL solution, 0.8 μmol), **3** in methanol (25 μL of a 6.7 mg/ mL solution, 0.4 μmol), and peptide in methanol (0.4 μmol) were added to a glass vial and diluted with methanol to a final volume of

500 μL. The mixture was vortexed for 30 min before the solvent was removed under vacuum. A micellar solution of surfactant **4** (9.3 mg, 20 μmol) in $H_2O$ (2.0 mL) was added to the above template–FM complex, followed by the addition of divinylbenzene (2.8 μL, 20 μmol) and photoinitiator 2,2-dimethoxy-2-phenylacetophenone in DMSO (10 μL of a 12.8 mg/mL solution, 0.5 μmol). The mixture was subjected to ultrasonication for 10 min before compound **5** (4.1 mg, 24 μmol), $CuCl_2$ in $H_2O$ (10 μL of 6.7 mg/mL solution, 0.5 μmol) and sodium ascorbate in $H_2O$ (10 μL of 99 mg/mL solution, 5 μmol) were added. The reaction mixture was stirred slowly at room temperature for 12 h. After incubation, compound **6** (10.6 mg, 40 μmol), $CuCl_2$ in $H_2O$ (10 μL of 6.7 mg/mL solution, 0.5 μmol), and sodium ascorbate in $H_2O$ (10 μL of 99 mg/mL solution, 5 μmol) were added. The mixture was stirred at room temperature for another 6 h, then purged with nitrogen for 15 min. The vial was sealed with a rubber stopper, degassed and refilled with nitrogen three times, and irradiated in a Rayonet reactor for 12 h. Finally, the reaction mixture was poured into acetone (8 mL). The precipitate was collected by centrifugation and washed with a mixture of acetone/water (5 mL/1 mL) three times, methanol/acetic acid (5 mL/0.1 mL) three times, acetone (5 mL) twice, and then dried in air to afford the final MINPs. Typical yields were > 80%.

**ITC Titration Experiments**

A solution of the titrating peptide was injected in equal steps of 10 μL into the sample cell containing the MINP solution. In the ITC curves presented, the top panel shows the raw calorimetric data. The area under each peak represents the amount of heat generated at each injection and is plotted against the molar ratio of the peptide to the MINP (Figure S7). The solid line is the best fit of the experimental data to the sequential binding of *N* binding sites on the MINP. The heat of dilution, obtained by adding the titrating solution to the buffer, was subtracted from the heat released during the binding. Binding parameters were generated from curve fitting using Microcal Origin 7.0.

**Protein Purification**

Protein constructs used in this study have been previously outlined[26]. K60P, K60A, E404P, and Y402F Pyk2 FERM−kinase mutants were generated using the QuikChange (Agilent Genomics) site-directed mutagenesis strategy. All variants were confirmed by DNA sequencing. Purification of all proteins used in this study was performed as previously described[26]. Briefly, Pyk2 FERM−kinase constructs (WT, K60P, K60A, E404P, and Y402F) or $His_6$-SUMO-Src constructs (WT [1-536] or ΔSH2 Src [1-142,166-536]) were co-transformed with the YopH phosphatase plasmid (pESU009), encoding tyrosine phosphatase domain (residues 177 – 463), into *E. coli* NiCo21(DE3) cells to generate homogenous, dephosphorylated Pyk2 or Src proteins. After overnight expression at 18°C, cells were harvested and lysed by sonication in lysis buffer composed of 50 mM Tris pH 8, 25 mM NaCl, 20 mM imidazole, 1 mM

PMSF, 5 mM EDTA, 5% glycerol, 2.5 mM β-mercaptoethanol, and protease inhibitors. Lysates were cleared by ultracentrifugation at 80,000 ×g for 15 min at 4 °C. $His_6$-tagged Pyk2 or $His_6$-tagged Src was isolated by Ni-NTA affinity chromatography, followed by cleavage of the $His_6$-SUMO tag using $His_6$-tagged Ulp1 protease. A subtractive passage through Ni-NTA resin removed the protease and cleaved $His_6$-SUMO tag. The purified Pyk2 or Src constructs were concentrated via centrifugal filtration and further purified by gel filtration chromatography in SEC buffer composed of 50 mM HEPES pH 7.4, 150 mM NaCl, 5% glycerol, 5 mM TCEP. Purified Pyk2 constructs or Src was concentrated, aliquoted, frozen in liquid nitrogen, and stored at -80 °C.

**Kinase Assays**

Kinase reactions were performed using a final concentration of 0.5 μM or 1 μM for wild type (WT) or Pyk2 FERM–kinase variants in a buffer consisting of 50 mM HEPES pH 7.4, 150 mM NaCl, 8 mM $MgCl_2$, 5% glycerol. The secondary structure in Pyk2 was investigated in the presence of MINP(Y402) or non-imprinted nanoparticle (NINP). MINP(Y402) was added to individual kinase reactions at a range of final concentrations (1, 3, or 6 μM), whereas NINP was added at 6 μM. Stock solutions were sonicated before use for 5min. Kinase/MINP mixtures were pre-equilibrated at 20 °C for 5 min. Kinase activity was initiated by adding 4 mM ATP (final concentration) and quenched at specific time points with 10 mM EDTA followed by heat denaturation at 90°C for 60 seconds.

To investigate the Pyk2–Src interaction, we performed kinase assays similar to the above protocol. Specifically, WT Pyk2 FERM−kinase (1 μM) was pre-incubated with kinase buffer containing 4 mM ATP for 10 minutes at 20 °C to ensure autophosphorylation at Y402 residue in Pyk2. WT Src OR ΔSH2 Src was subsequently added to the reaction mixture at a final concentration of 0.5 μM to monitor Pyk2 activation loop phosphorylation. Samples were quenched at various time points with 10 mM EDTA (final concentration) followed by heat denaturation at 90°C for 60 seconds.

To study the ability of MINPs to block Src binding motif. We performed kinase assays in which WT Pyk2 FERM−kinase (1 μM) was pre-incubated with kinase buffer containing 4 mM ATP for 20 minutes at 20 °C to ensure autophosphorylation at Y402 residue in Pyk2. MINPs (pY402) and (PRR) were subsequently added to the reaction mixture at a final concentration of 6 μM and incubated for 3 minutes. Src was then added to the reaction to monitor the phosphorylation of Y579/Y580 in Pyk2. Samples were quenched following 5 min of Src addition or at various time points with 10 mM EDTA (final concentration) followed by heat denaturation at 90°C for 60 seconds.

General phosphotyrosine production was assessed via Western or Dot blotting using primary mouse anti-phosphotyrosine antibody (PY20) followed by goat-anti-mouse HRP conjugated secondary antibody. Site-specific phosphotyrosine blotting was performed with anti-phospho-PTK2B (pY402) (Sigma-Aldrich, RRID: AB_10624208) or anti-phospho-PYK2 (pY579/pY580) (Invitrogen, RRID: AB_2533706) primary antibodies with the goat anti-mouse HRP-conjugated secondary antibody. Blots were imaged with a chemiluminescent substrate (Pierce). Phosphotyrosine density was quantified by densitometry (Image Lab, Bio-Rad).

**Global Hydrogen/Deuterium Exchange-Mass Spectrometry (HDX-MS)**

Global HDX-MS was initiated by diluting Pyk2 FERM-kinase variants (80 pmol) into $D_2O$ buffer composed of a final concentration of 150 mM KCl, 50 mM HEPES, 5 mM DTT (pD 7.4), and 89% $D_2O$. Exchange reaction mixtures were incubated at 21 °C. At time point 2.5 min, labeling reactions were quenched to pH 2.5 by addition of 0.25M sodium phosphate (final concentration). Non-exchanged samples were acquired in matched $H_2O$ buffers to establish intact mass baselines for each variant. Quenched samples were immediately injected (2 µL) into an ACQUITY UPLC H-class instrument coupled in line to an ESI-Q-TOF Synapt G2-Si instrument (Waters). Mobile phases consisted of solvents A (HPLC-grade aqueous 0.1% formic acid) and B (HPLC-grade acetonitrile and 0.1% formic acid). Exchange samples were desalted on a C4 column (5 µm, 1 mm × 50 mm; Restek) with 30% solvent B for 1 min at a flow rate of 400 µL/min. Intact protein was eluted with a rapid gradient ramp from 35% to 75% solvent B over 3 min. MS data were collected in positive ion, MS continuum, and resolution mode with an *m*/*z* range of 300–4000. The intact mass of each exchange time point was deconvoluted from multiple charge states using the MaxEnt1 function of MassLynx (Waters).

**Quantification and statistical analysis**

The information regarding quantification and statistical analysis is indicated in the figure legends. For kinase assays, n = 3-5 where n is the number of replicate assays/measurements. Statistical significance was determined using a two-tailed unpaired t test. The individual values were plotted with the mean and standard deviation on GraphPad Prism 9.

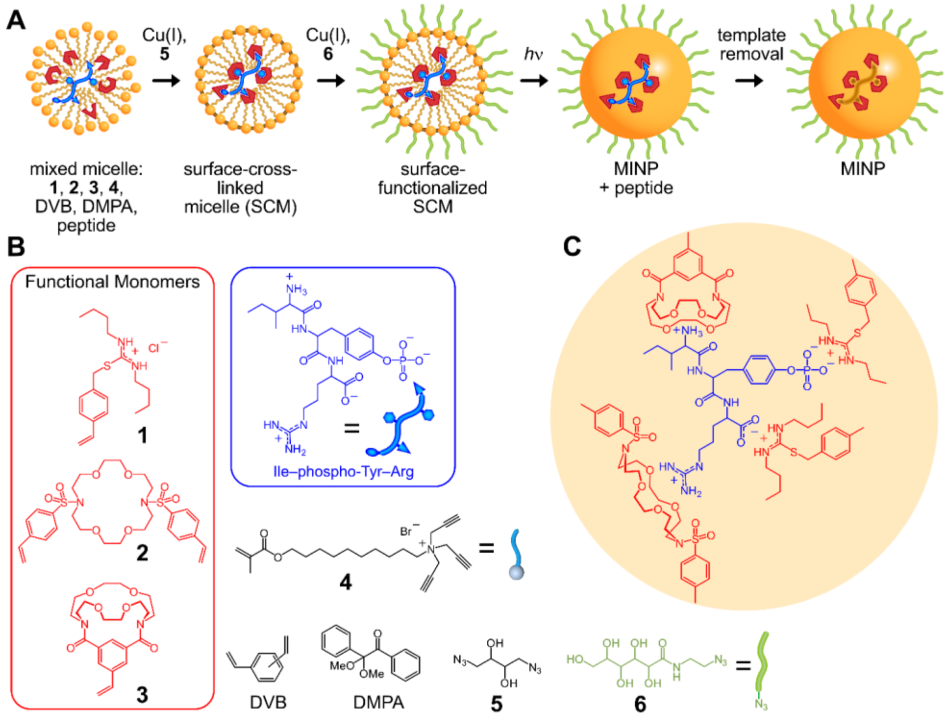


**Scheme S1. Synthesis of peptide-templated MINP.** (A) Peptide embeds with mixed micelles of functional monomers **1**, **2**, and **3** with cross-linkable surfactant **4** to provide specific, complementary hydrophobic, charge, and hydrogen bonding interactions. Mixed micelle surfaces are cross-linked via Cu(I)-catalyzed azide-alkyne cycloaddition with **5,** followed by surface functionalization with **6**. Core cross-linking to generate imprinted MINP proceeds via radical polymerization by activating (*h*ν) photoinitiator DMPA. (B) Structures of functional monomers (**1**, **2**, **3**), micellar surfactant (**4**), cross-linking reagents (**5**, divinylbenzene, DVB; 2,2-dimethoxy-2-phenylacetophenone, DMPA), surface solubilization moiety (**6**), and an example phosphopeptide. (C) Illustration of functional monomer interactions with an example peptide. Functional monomer **1** confers specific interactions with carboxylates and phosphates, **2** with arginine guanidiniums, and **3** for amines. Adapted from Li, et al[8].

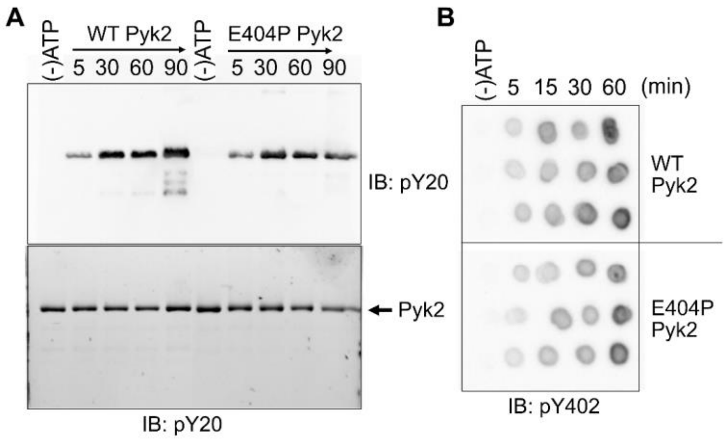


**Fig. S1. Kinase activity time course of WT and E404P Pyk2 FERM−kinase.** Autophosphorylation activity of Pyk2 FERM–kinase variants (0.5 μM) was initiated by adding ATP (4 mM) at 21 °C. Time points were quenched with 10 mM EDTA and probed by immunoblotting with (a) pan-specific anti-phosphotyrosine primary antibody (PY20) or (b) anti-phospho-PTK2B (pY402). Protein loading was imaged by 2,2,2-trichloroethanol staining. In (b) each row of dots represents one assay replicate time course.

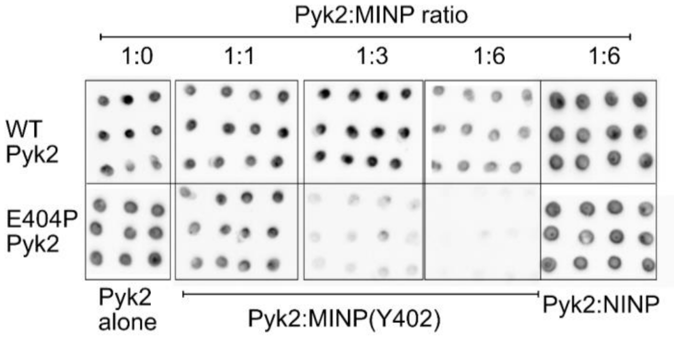


**Fig. S2. Pyk2 autophosphorylation inhibition by MINP(Y402).** WT or E404P Pyk2 FERM–kinase (0.5 μM) was incubated with MINP(Y402) at a range of final concentrations (1, 3, or 6 μM) for 5 min at 21 °C. NINP was tested at 6 μM in the same conditions as MINP(Y402). Autophosphorylation (phospho-Y402) of WT or E404P Pyk2 FERM–kinase was initiated by addition of 4 mM ATP and quenched with 100 mM EDTA. Samples were probed by immunoblotting with pan-specific anti-phosphotyrosine primary antibody (PY20). Each column of dots represents one assay replicate.

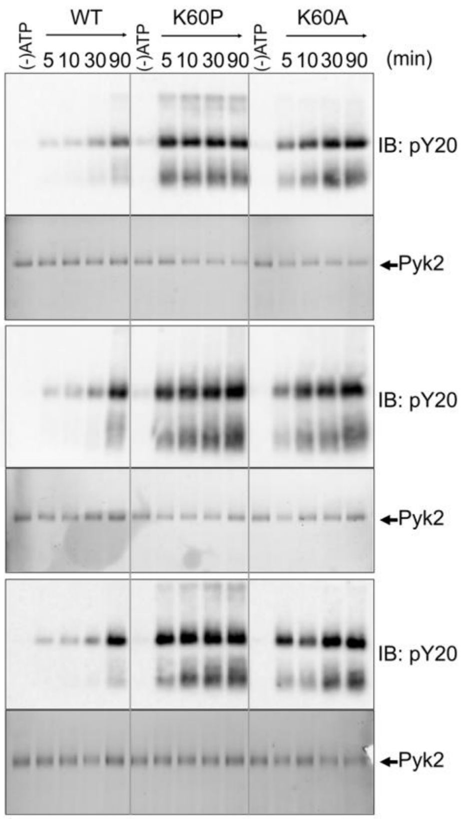


**Fig. S3. FERM domain variants modulate Pyk2 autophosphorylation.** Autophosphorylation activity time course of WT, K60A, and K60P Pyk2 FERM–kinase variants (0.5 μM) was initiated by adding ATP (4 mM) at 22 °C. Time points were quenched with 10 mM EDTA. Phosphorylation of Pyk2 variants at residue Y402 was measured by Western blotting using pan-specific anti-phosphotyrosine primary antibody (PY20) primary antibody. Protein loading (bottom rows) was imaged by 2,2,2-trichloroethanol staining and used to normalize the densitometry data from the western blots (top rows). Experiment was performed in triplicate.

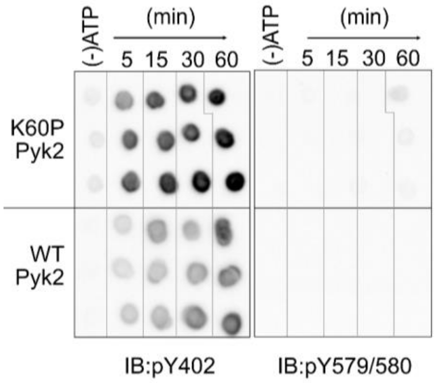


**Fig. S4. The autophosphorylation target of K60P Pyk2.** Autophosphorylation activity time course of WT and K60P Pyk2 FERM–kinase variants (0.5 µM) was initiated by adding ATP (4 mM) at 22 °C and quenched with 10 mM EDTA. Site-specific phosphorylation was detected by blotting with anti-phospho- PYK2 (pY579/Y580) or anti-phospho-PTK2B (pY402). Each row of dots represents one assay replicate time course.

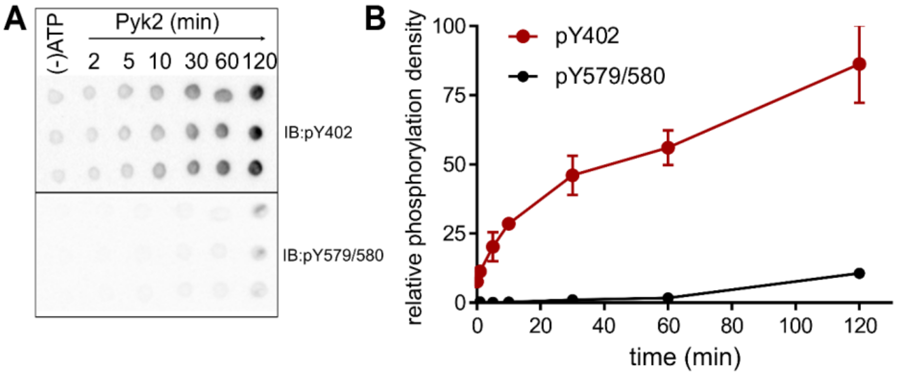


**Fig. S5. Role of Src kinase in Pyk2 phosphorylation.** (A) To investigate the ability of Pyk2 to phosphorylate its own activation loop, the phosphorylation of Pyk2 was initiated by addition of 4 mM ATP and monitored for over 60 min at 21 °C. Time points were quenched with 100 mM EDTA and probed by immunoblotting with anti-phospho-pY579/pY580 or anti-phospho-Y402 primary antibody. Each row of dots represents one assay replicate time course. (B) Quantification of (A) by densitometry.

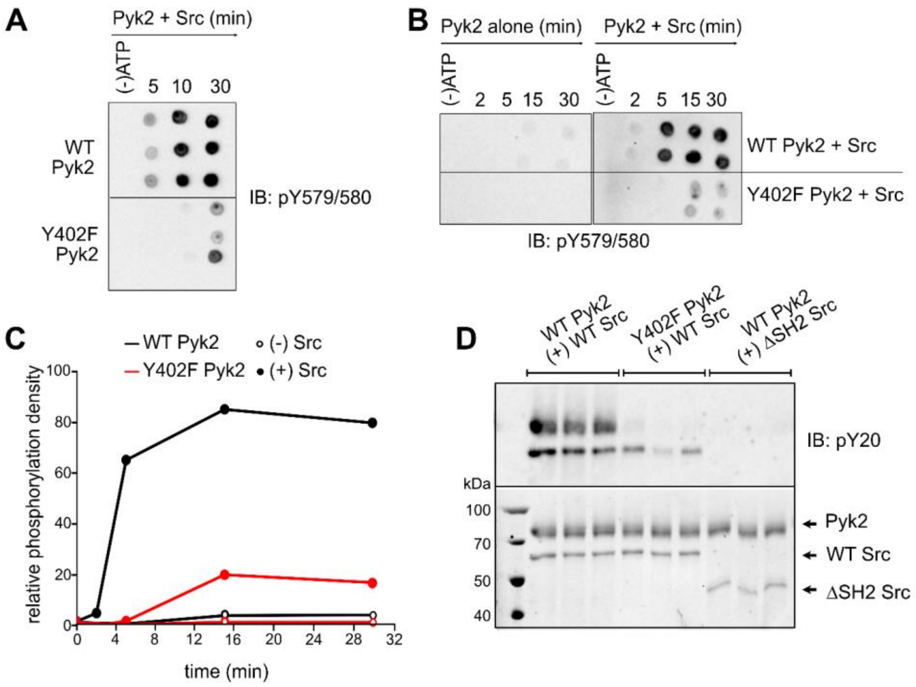


**Fig. S6. Pyk2 phosphorylation in the presence of Src.** (A) The Src-mediated phosphorylation time course of WT or Y402F Pyk2 FERM–kinase was initiated by adding 4 mM ATP. Reaction time points were sampled over 30 min at 21 °C. For both (A) and (B) time points were quenched with 10 mM EDTA and probed by immunoblotting with anti-phospho-PYK2 (pY579/Y580) primary antibody. Each row of dots represents one assay replicate time course. (C) Quantification of (B) by densitometry. (D) Activity time course was measured for WT Pyk2 or Y402F Pyk2 (1µM) at 21 °C. Pyk2 was incubated with 4mM ATP for 10 min for autophosphorylation before addition of 0.5 µM of WT Src or ΔSH2 Src [1-142,166-536]. Samples were quenched with EDTA 10 min following the addition of Src.

**Commented [EU1]:** Did we forget Y402F here?

**Commented [EU2R1]:** Currently, we're relying on the gel bands into indicate which reactions contain which Src construct. Let's help the reader by adding labels about the Pyk2 constructs indicating which Src was included (like in Fig. 4).

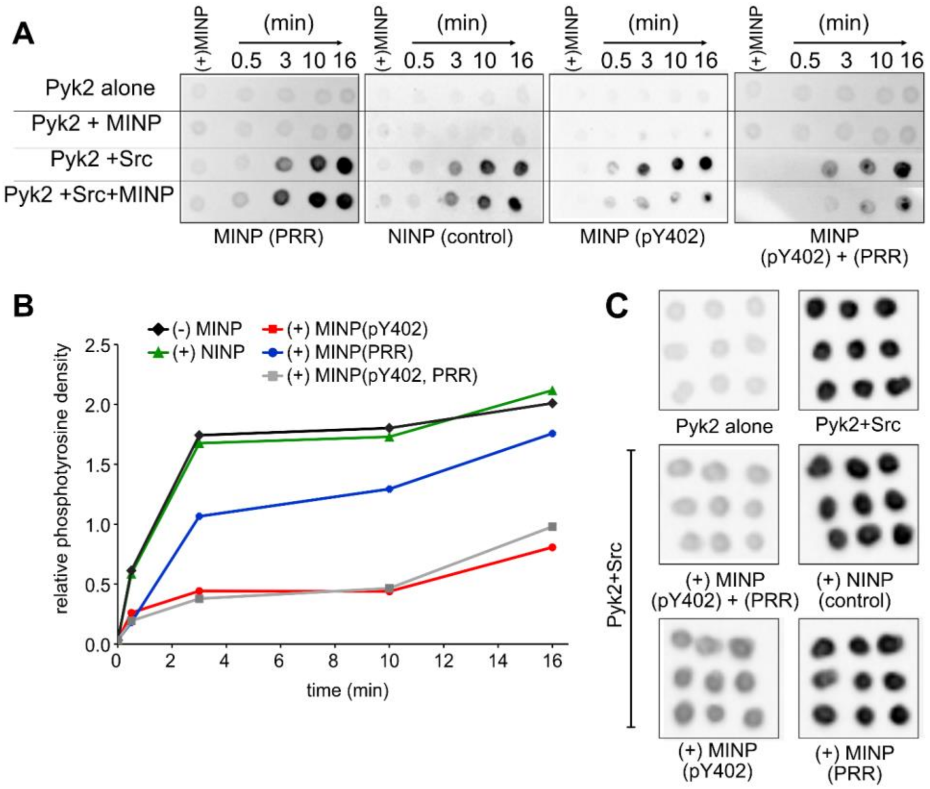


**Fig. S7. Interaction sites in the Pyk2−Src complex probed by MINPs.** (A) and (C) Kinase activity of Pyk2 FERM-kinase (1 μM) was assessed in the presence or absence of Src (0.5 μM). Phosphorylation of the Y402 site on Pyk2 FERM−kinase was measured after a 3-minute incubation with MINP, followed by the addition of Src. (B) Phosphorylation levels of Pyk2 Y579/Y580 at different time points (A) were quantified using densitometry analysis. (C) Data collected at the 5-minute time point were subjected to site-specific phosphorylation detection by blotting with anti-phospho-PYK2 (pY579/Y580). Each column of dots in (C) represents one assay replicate.

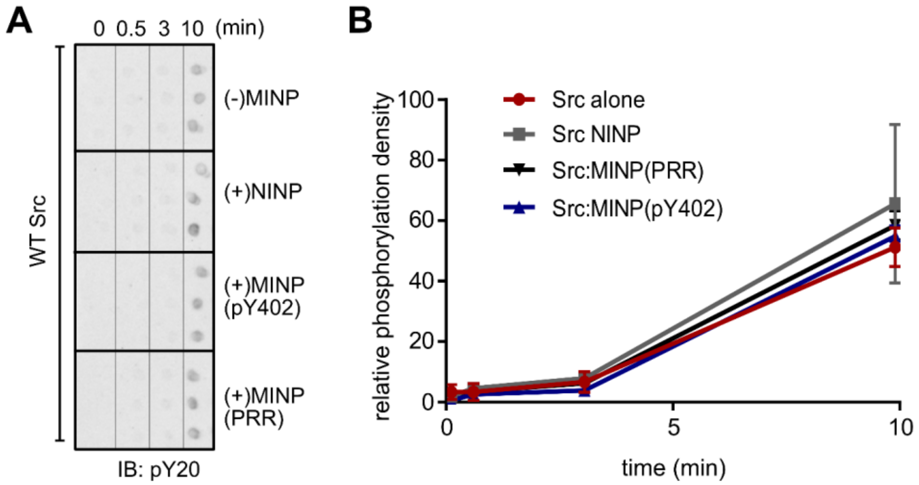


**Fig. S8. MINPs have no inhibitory effect towards Src.** (A) Kinase activity of Src (1 µM) was assessed in the presence or absence of MINPs or NINP (6 µM). Autophosphorylation of the Src was measured after a 3-minute incubation with MINP, followed by the addition of ATP (4 mM). Phosphorylation levels of Src at different time points were detected by blotting with pan-specific anti-phosphotyrosine primary antibody (PY20) and (B) quantified using densitometry. Each row of dots in (A) represents one activity assay replicate.

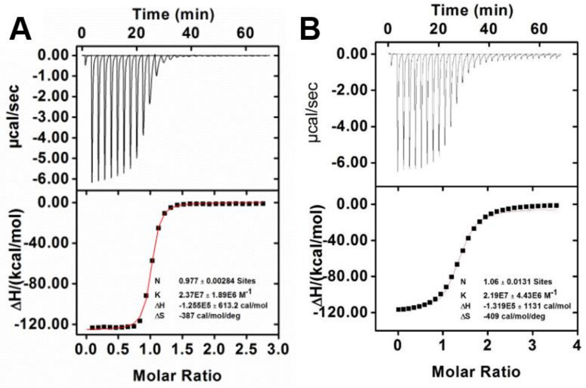


**Fig. S9. Binding affinity of (A) MINP(PRR) and (B) MINP(pY402) towards peptide derived from Pyk2 sequence.** ITC titration curves were obtained at 298 K for the titration of the (A) peptide PRR corresponding to the Pyk2 proline-rich sequence (RNSLPQIPMLN) and for (B) phosphorylated peptide corresponding to Pyk2 pY402 sequence (DI**pY**AEIPDETLR) with MINPs prepared with FMs. More details about the titration curve used in this study can be found in Li, et al[17].
